# YAP/TAZ inactivation with simvastatin attenuates glucocorticoid-induced human trabecular meshwork cell dysfunction

**DOI:** 10.1101/2022.09.27.509788

**Authors:** Hannah Yoo, Ayushi Singh, Haiyan Li, Ana N. Strat, Tyler Bagué, Preethi S. Ganapathy, Samuel Herberg

## Abstract

**Purpose:** Impairment of the trabecular meshwork (TM) is the principal cause of increased outflow resistance in the glaucomatous eye. Yes-associated protein (YAP) and transcriptional coactivator with PDZ binding motif (TAZ) are emerging as potential mediators of TM cell/tissue dysfunction. Furthermore, YAP/TAZ activity was recently found to be controlled by the mevalonate pathway in non-ocular cells. Clinically-used statins block the mevalonate cascade and were shown to improve TM cell pathobiology; yet, the link to YAP/TAZ signaling was not investigated. In this study, we hypothesized that YAP/TAZ inactivation with simvastatin attenuates glucocorticoid-induced human TM (HTM) cell dysfunction.

**Methods:** Primary HTM cells were seeded atop or encapsulated within bioengineered extracellular matrix (ECM) hydrogels. Dexamethasone was used to induce a pathologic phenotype in HTM cells in the absence or presence of simvastatin. Changes in YAP/TAZ activity, actin cytoskeletal organization, phospho-myosin light chain levels, hydrogel contraction/stiffness, and fibronectin deposition were assessed.

**Results:** Simvastatin potently blocked pathologic YAP/TAZ nuclear localization/activity, actin stress fiber formation, and myosin light chain phosphorylation in HTM cells. Importantly, simvastatin co-treatment significantly attenuated dexamethasone-induced ECM contraction/stiffening and extracellular fibronectin deposition. Sequential treatment was similarly effective but did not match clinically-used Rho kinase inhibition.

**Conclusions:** YAP/TAZ inactivation with simvastatin attenuates HTM cell pathobiology in a tissue-mimetic ECM microenvironment. Our data may help explain the association of statin use with a reduced risk of developing glaucoma via indirect YAP/TAZ inhibition as a proposed regulatory mechanism.

## Introduction

The trabecular meshwork (TM) plays a central role in the conventional outflow pathway, which drains the aqueous humor from the anterior chamber to regulate outflow facility and intraocular pressure^1–3^. The bidirectional interactions between TM cells^4^ and their extracellular matrix (ECM) are crucial for maintaining normal tissue function in the healthy eye^5,6^. In primary open-angle glaucoma, the most common form of glaucoma^7^, disruption of these interactions drives progressive fibrotic-like tissue remodeling. Key characteristics of this process include increased TM contraction, actin stress fiber assembly, ECM deposition/crosslinking, and overall tissue stiffening^8^. These pathologic alterations lead to increased outflow resistance driving ocular hypertension, which provides further negative feedback and may ultimately push the TM to irreversibly fail^9,10^. Despite substantial scientific effort over the past several decades devoted to understanding TM pathophysiology, the mechanisms underlying persistent tissue dysfunction in glaucoma remain elusive.

Yes-associated protein (YAP) and transcriptional coactivator with PDZ-binding motif (TAZ) are the downstream mediators of the Hippo pathway that play important roles in tissue homeostasis and organ growth^11,12^. As mechanotransducers, activated YAP/TAZ translate biophysical stresses into biochemical signals by translocating to the nucleus from the cytoplasm and interacting with TEAD transcription factors. Through this mechanism, YAP/TAZ exert their function on regulating cellular gene expression, proliferation, and fate^13^. Imbalance or failure of this process is central to a variety of disorders^14^ YAP/TAZ hyperactivity is strongly associated with glaucomatous TM cell dysfunction. Our group^15^ and others showed that various glaucoma-associated stressors (e.g., ECM stiffness, dexamethasone, transforming growth factor ß2) increase YAP/TAZ activity independent of the canonical Hippo pathway^16–22^. Importantly, *YAP1* was recently identified as one of forty-four previously unknown open-angle glaucoma-risk loci across European, Asian, and African ancestries^23^, suggesting a potential causal relationship with TM outflow dysfunction. Therefore, targeting YAP/TAZ signaling - directly or indirectly - may be of therapeutic value for treating outflow dysfunction in primary open-angle glaucoma.

Statins are widely used cholesterol-lowering oral medications for preventing and treating cardiovascular diseases^24^ They block the conversion of hydroxymethylglutaryl coenzyme A (HMG-CoA) to mevalonate by competitively inhibiting HMG-CoA reductase, the rate-limiting enzyme of the mevalonate pathway^25^. Emerging clinical reports suggest that ocular hypertensive/glaucoma patients have higher total cholesterol levels than patients without glaucoma^26,27^ Statin use has also been associated with a reduced risk of primary open-angle glaucoma development and progression in some studies^28–33^, possibly contributed to by statins’ pleiotropic effects including anti-inflammatory and anti-oxidative benefits. Yet, mechanistic details underlying this potential protective effect in relation to tissues/cells of the outflow tract are incompletely understood.

The ability of statins to induce TM cell relaxation, as evidenced by changes in the actin cytoskeleton, and to increase aqueous humor outflow facility in porcine eye anterior segments was initially demonstrated using lovastatin^34^ By the same token, lovastatin was shown to cause marked changes in human TM cell morphology including a loss of filamentous (F)-actin stress fiber organization^35^, concomitant with a marked accumulation of cytosolic inactive Rho GTPase proteins^36^. Atorvastatin, another lipophilic statin with higher potency compared to lovastatin^37^, was found to reduce ECM protein expression in human TM cells^38^, as well as induce significant changes in cellular morphology and focal adhesions^39^. Importantly, it was shown that YAP/TAZ activity is controlled by the mevalonate pathway. In two independent small-scale library screens, statins including simvastatin and lovastatin were found to elicit strong YAP/TAZ inhibitory effects potently opposing nuclear localization and transcriptional responses in a Rho GTPase-dependent manner^40,41^. These observations suggest that statins may exert their beneficial clinical effects through modulating metabolic processes independent from cholesterol homeostasis. Therefore, targeting YAP/TAZ signaling with statins presents an intriguing avenue for mitigating TM cell pathobiology in glaucoma with translational potential.

Clinically, glucocorticoid exposure can result in ocular hypertension and may lead to the development of steroid-induced glaucoma, which has commonalities with primary open-angle glaucoma^42,43^. Treatment of TM cells with the synthetic glucocorticoid dexamethasone likewise induces cells to undergo a pathological phenotypic conversion^44,45^. Therefore, dexamethasone is widely used to reliably induce glaucoma-like TM cell pathobiology *in vitro*. In this study, we hypothesized that YAP/TAZ inactivation with simvastatin attenuates dexamethasone-induced human TM cell dysfunction - assessed by quantifying alterations in YAP/TAZ sub-cellular localization/activity, actomyosin cytoskeletal organization, and functional ECM contraction/remodeling - in a soft tissue-mimetic 3D ECM hydrogel^46^.

## Materials and Methods

### HTM cell isolation and culture

The use of human donor corneas was approved by the SUNY Upstate Medical University Institutional Review Board (protocol #1211036), and all experiments were performed according to the tenets of the Declaration of Helsinki for the use of human tissue. Primary human TM (HTM) cells were isolated from healthy donor corneal rims discarded after transplant surgery, as previously described^15,46–48^, and cultured according to established protocols^49,50^. Five normal, previously characterized HTM cell strains (HTM05, HTM12, HTM14, HTM19, HTM36) were used in this study (**Table. 1**). All HTM cell strains were validated with dexamethasone-induced (100 nM) myocilin expression in more than 50% of cells by immunocytochemistry and immunoblot analyses. Different combinations of 2-3 HTM cell strains were used per experiment, depending on cell availability, and all studies were conducted between cell passage 3-7. HTM cells were cultured in low-glucose Dulbecco’s Modified Eagle’s Medium (DMEM; Gibco; Thermo Fisher Scientific, Waltham, MA, USA) containing 10% fetal bovine serum (FBS; Atlanta Biologicals, Flowery Branch, GA, USA) and 1% penicillin/streptomycin/glutamine (PSG; Gibco) and maintained at 37°C in a humidified atmosphere with 5% CO_2_. Fresh media was supplied every 2-3 days.

**Table 1.**
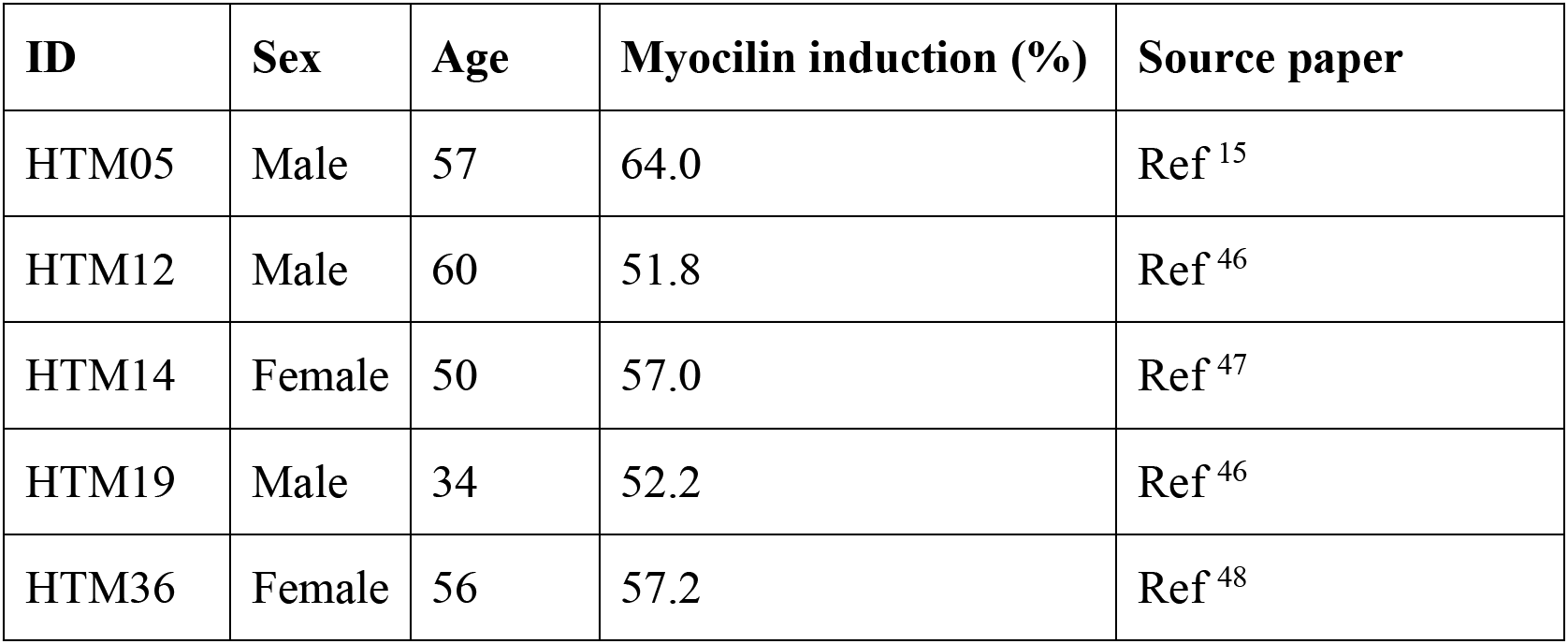
HTM cell strain information.

### Hydrogel precursor solutions

Methacrylate-conjugated bovine collagen type I (MA-COL; Advanced BioMatrix, Carlsbad, CA, USA) was reconstituted in sterile 20 mM acetic acid to achieve 6 mg/ml. Immediately prior to use, 1 ml MA-COL was neutralized with 85 μl neutralization buffer (Advanced BioMatrix) according to the manufacturer’s instructions. Thiol-conjugated hyaluronic acid (SH-HA; Glycosil^®^; Advanced BioMatrix) was reconstituted in sterile diH2O containing 0.5% (w/v) photoinitiator (4-(2-hydroxyethoxy) phenyl-(2-propyl) ketone; Irgacure^®^ 2959; Sigma-Aldrich) to achieve 10 mg/ml according to the manufacturer’s protocol. In-house expressed ELP (SH-ELP; thiol via KCTS flanks^46,51^) was reconstituted in chilled DPBS to achieve 10 mg/ml and sterilized using a 0.2 μm syringe filter in the cold.

### HTM hydrogel preparation

Hydrogel precursors MA-COL (3.6 mg/ml [all final concentrations]), SH-HA (0.5 mg/ml, 0.025% (w/v) photoinitiator), and SH-ELP (2.5 mg/ml) were thoroughly mixed in an amber color tube on ice. Thirty microliters of the hydrogel solution were pipetted onto Surfasil-coated (Fisher Scientific) 18 × 18-mm square glass coverslips followed by placing 12-mm round glass coverslips onto the hydrogels to facilitate even spreading of the polymer solution. Hydrogels were crosslinked by exposure to UV light (OmniCure S1500 UV Spot Curing System; Excelitas Technologies, Mississauga, Ontario, Canada) at 320-500 nm, 2.2 W/cm^2^ for 5 s, according to our established protocols^15,46–48^. The coverslips were removed with fine-tipped tweezers and placed hydrogel-side facing up in polydimethylsiloxane-coated (PDMS; Sylgard 184; Dow Corning; Fisher Scientific) 24-well culture plates before seeding HTM cells (1.5 × 10^4^ cells/cm^2^) atop. To fabricate larger ECM hydrogels for immunoblot analyses, 500 μl of the hydrogel solution were pipetted into 6-well culture plates for uniform coverage across the entire well surface and crosslinked using a modified UV protocol (80 mW/cm^2^ for 30 s) before seeding HTM cells (1.5 × 10^5^ cells/cm^2^) atop. These adjusted settings were shown to yield ECM hydrogels with equivalent elastic modulus compared to standard hydrogels (**Suppl. Fig. 1**). For cell encapsulated hydrogels, HTM cells (1.0 × 10^6^ cells/ml) were thoroughly mixed with the hydrogel precursors on ice, followed by pipetting either 10 μl droplets of the mixture onto PDMS-coated 24-well culture plates, or 250 μl into custom 16 × 1-mm PDMS molds and routine photocrosslinking.

### HTM hydrogel treatments

HTM cells seeded atop ECM hydrogels were cultured in DMEM with 10% FBS and 1% PSG for 1-3 days until ~80-90% confluent. Then, constructs were cultured in serum-free DMEM with 1% PSG and subjected to the following treatments for 3 days: **1) control** (vehicle: 0.1% ethanol; 0.1% dimethyl sulfoxide (DMSO), both from Fisher Scientific), **2) dexamethasone** (100 nM in ethanol; Fisher Scientific), **3) simvastatin** (10 μM in DMSO; Sigma-Aldrich, St. Louis, MO, USA), **4) dexamethasone + simvastatin** (100 nM dexamethasone; 10 μM simvastatin), and **5) dexamethasone + simvastatin + mevalonate-5-phosphate** (100 nM dexamethasone; 10 μM simvastatin; 500 μM mevalonate-5-phosphate in water; Sigma-Aldrich). HTM cell-encapsulated ECM hydrogels were cultured in DMEM with 10% FBS and 1% PSG and subjected to the same treatments for 10 days. The co-treatment strategy was chosen to simulate a “prophylactic treatment approach”.

In another set of experiments, HTM hydrogels were first induced with dexamethasone for 5 days followed by different rescue treatments for 5 days with dexamethasone withheld. This sequential treatment strategy was chosen to simulate a “therapeutic treatment approach”. HTM cell-encapsulated ECM hydrogels were cultured in DMEM with 10% FBS and 1% PSG and subjected to the following treatments for 10 days: **1) control** (vehicle: 0.1% ethanol [0-5 d]; 0.1% DMSO [5-10 d]), **2) dexamethasones** (100 nM dexamethasone [0-5 d]; 0.1% DMSO [5-10 d]), **3) dexamethasones + simvastatins** (100 nM dexamethasone [0-5 d]; 10 μM simvastatin [5-10 d]), **4) dexamethasones + netarsudils** (100 nM dexamethasone [0-5 d]; 1.0 μM netarsudil in DMSO;

Aerie Pharmaceuticals, Durham, NC, USA [5-10 d]). The 5-day dexamethasone exposure was shown to result in equivalently-induced HTM hydrogel contraction compared to the standard 10 days (**Suppl. Fig. 2**). The dexamethasone concentration was selected based on our previous study^46^. The simvastatin and mevalonate-5-phosphate concentrations were selected according to a previous report^40^. The netarsudil concentration was selected based on our recent study^48^.

### HTM hydrogel immunocytochemistry analysis

HTM cells cultured atop ECM hydrogels in presence of the different treatments for 3 days were fixed with 4% paraformaldehyde (Thermo Fisher Scientific) at room temperature for 20 min, permeabilized with 0.5% Triton™ X-100 (Thermo Fisher Scientific), blocked with blocking buffer (BioGeneX, Fremont, CA, USA), and incubated with primary antibodies, followed by incubation with fluorescent secondary antibodies (**Table 2**); nuclei were counterstained with 4’,6’-diamidino-2-phenylindole (DAPI; Abcam, Waltham, MA, USA). Similarly, cells were stained with DyLight™ 594 Phalloidin (Cell Signaling Technology, Danvers, MA, USA)/DAPI (Abcam) according to the manufacturer’s instructions. Coverslips were mounted with ProLong™ Gold Antifade (Invitrogen; Thermo Fisher Scientific) on Superfrost™ microscope slides (Fisher Scientific), and fluorescent images were acquired with an Eclipse N*i* microscope (Nikon Instruments, Melville, NY, USA).

**Table 2.**
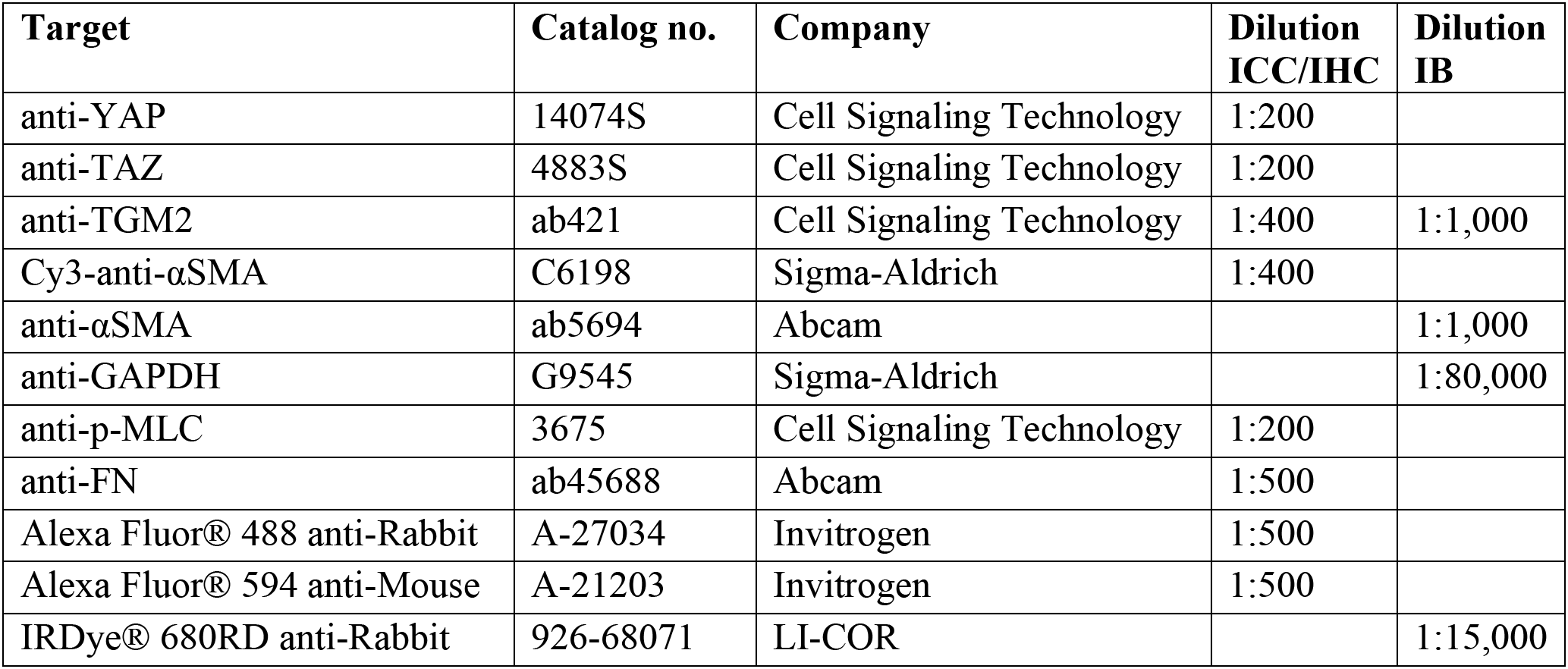
Antibody information.

All fluorescent image analyses were performed using FIJI software (National Institutes of Health (NIH), Bethesda, MD, USA)^52^. The cytoplasmic YAP/TAZ intensity was measured by subtracting the overlapping nuclear (DAPI) intensity from the total YAP/TAZ intensity. The proportion of total YAP/TAZ intensity that overlapped with the nucleus (DAPI) was obtained to measure the nuclear YAP/TAZ intensity. YAP/TAZ nuclear/cytoplasmic (N/C) ratio was calculated as follows: N/C ratio = (nuclear signal/area of nucleus) / (cytoplasmic signal/area of cytoplasm). YAP/TAZ N/C ratios and fluorescence signal intensities of TGM2, F-actin, αSMA, and p-MLC were measured in at least N = 9 images from 3 experimental replicates per treatment group from 3 HTM cell strains with image background subtraction, followed by calculation of fold-change vs. control.

### HTM hydrogel immunoblot analysis

HTM cells cultured atop ECM hydrogels in presence of the different treatments for 3 days were lifted from the hydrogel using 0.25% trypsin/EDTA (Gibco; Fisher Scientific). Care was taken not to contaminate the cellular fraction with ECM proteins from the hydrogel substrate. Pooled cells from 3 experimental replicates per group were lysed in lysis buffer (CelLytic™ M, Sigma-Aldrich) supplemented with Halt™ protease/phosphatase inhibitor cocktail (Thermo Fisher Scientific). Equal protein amounts (10 μg), determined by standard bicinchoninic acid assay (Pierce; ThermoFisher Scientific), in 4× loading buffer (Invitrogen; Thermo Fisher Scientific) with 5% beta-mercaptoethanol (Fisher Scientific) were boiled for 5 min and subjected to SDS-PAGE using NuPAGE™ 4-12% Bis-Tris Gels (Invitrogen; Thermo Fisher Scientific) at 150V for 120 min and transferred to 0.45 μm PVDF membranes (Sigma; Thermo Fisher Scientific). Membranes were blocked with 5% bovine serum albumin (Thermo Fisher Scientific) in tris-buffered saline with 0.2% Tween^®^20 (Thermo Fisher Scientific), and probed with primary antibodies followed by incubation with fluorescent secondary antibodies (**Table 2**). Bound antibodies were visualized with an Odyssey CLx imager (LI-COR, Lincoln, NE, USA).

### HTM hydrogel contraction analysis

Longitudinal brightfield images of HTM hydrogels subjected to the different treatments for 10 days were acquired at day 0 and day 10 with an Eclipse T*i* microscope (Nikon). Construct areas from N = 4-8 experimental replicates per treatment group from 3 HTM cell strains were quantified using FIJI software (NIH) and normalized to 0 d followed by normalization to controls.

### HTM hydrogel cell viability analysis

The number of viable cells inside HTM hydrogels subjected to the different treatments for 10 days was quantified with the CellTiter 96^®^ Aqueous Non-Radioactive Cell Proliferation Assay (Promega; Thermo Fisher Scientific) following the manufacturer’s protocol. HTM hydrogels were incubated with the staining solution (38 μl MTS, 2 μl PMS solution, 200 μl DMEM) at 37°C for 1.5 h. Absorbance at 490 nm was recorded using a spectrophotometer plate reader (BioTek, Winooski, VT, USA). Blank (DMEM with the staining solution)-subtracted absorbance values served as a direct measure of HTM cell viability from N = 4 experimental replicates per treatment group from 1 representative HTM cell strain.

### HTM hydrogel rheology analysis

HTM hydrogels subjected to the different treatments for 10 days were cut to size using an 8-mm diameter tissue punch. A Kinexus rheometer (Malvern Panalytical, Westborough, MA, USA) fitted with an 8-mm diameter parallel plate was used to measure hydrogel viscoelasticity. To ensure standard conditions across all experiments, the geometry was lowered into the hydrogels until a calibration normal force of 0.02 N was achieved. An oscillatory shear-strain sweep test (0.1-60%, 1.0 Hz, 25°C) was then applied to determine storage modulus (G’) and loss modulus (G”) in the linear region from N = 3-4 experimental replicates per treatment group from 2 HTM cell strains. Young’s modulus was calculated with E = 2 * (1 + v) * G’, where a Poisson’s ratio (v) of 0.5 for the ECM hydrogels was assumed^53^.

### HTM hydrogel immunohistochemistry analysis

HTM hydrogels subjected to the different treatments for 10 days were fixed in 4% paraformaldehyde (Fisher Scientific) at 4°C overnight, incubated in 30% sucrose (Fisher Scientific) at 4°C for 24 h, embedded in Tissue-Plus™ O.C.T. Compound (Fisher Scientific), and flash frozen in liquid nitrogen. Twenty micrometer cryosections were cut using a cryostat (Leica Biosystems Inc., Buffalo Grove, IL, USA) and collected on Superfrost™ Plus microscope slides (Fisher Scientific). Sections were permeabilized with 0.5% Triton™ X-100, blocked with blocking buffer (BioGeneX) and incubated with a primary antibody against fibronectin, followed by incubation with a fluorescent secondary antibody (**Table 2**). Slides were mounted with ProLong™ Gold Antifade (Thermo Fisher Scientific), and fluorescent images were acquired with an Eclipse N*i* microscope (Nikon). Fluorescence signal intensity of fibronectin was measured using FIJI (NIH) in at least N = 9 images from 4 experimental replicates per treatment group from 2 HTM cell strains with image background subtraction, followed by calculation of fold-change vs. control.

### Statistical analysis

Individual sample sizes are specified in each figure caption. Comparisons between groups were assessed by unpaired *t* test, one-way or two-way analysis of variance (ANOVA) with Tukey’s multiple comparisons *post hoc* tests, as appropriate. All data are shown with mean ± SD, some with individual data points. The significance level was set at p<0.05 or lower. GraphPad Prism software v9.3 (GraphPad Software, La Jolla, CA, USA) was used for all analyses.

## Results

### Simvastatin reduces pathologic YAP/TAZ nuclear localization and TGM2 levels in HTM cells

The transcriptional coactivators YAP and TAZ dynamically shuttle between the cytoplasm and nucleus to regulate gene expression; increased nuclear YAP/TAZ localization principally indicates enhanced transcriptional activity^54^ Our recent studies on human TM and Schlemm’s canal cells support this notion in ocular cells^15,55^. Moreover, the mevalonate pathway was shown to promote YAP/TAZ nuclear accumulation and activity in non-ocular cells^40^; therefore, we hypothesized that simvastatin decreases glucocorticoid-induced pathologic YAP/TAZ nuclear localization and signaling in HTM cells. Exposure to the dexamethasone significantly increased YAP and TAZ nuclear-to-cytoplasmic (N/C) ratios in HTM cells cultured atop ECM hydrogels (YAP: ~3.6; TAZ: ~4.2) compared to controls (YAP: ~2.2; TAZ: ~1.8) (**Fig. 1A-D**). These results were consistent with our previous study using TGFß2 to induce glaucomatous HTM cell dysfunction^15^. With simvastatin alone, we observed YAP/TAZ N/C ratios (YAP: ~2.3; TAZ: ~2.5) comparable to the control group. Importantly, co-treatment with dexamethasone + simvastatin significantly decreased YAP/TAZ N/C ratios (YAP and TAZ: ~2.2) compared to dexamethasone-induced HTM cells, restoring control levels. When mevalonate-5-phosphate was supplemented to bypass the simvastatin-mediated HMG-CoA reductase inhibition, we observed significantly increased YAP/TAZ N/C ratios (YAP: ~3.6; TAZ: ~3.5) compared to dexamethasone + simvastatin, indistinguishable from the dexamethasone-treated HTM cells. Different HTM cell strains significantly affected the results (p<0.0001), in agreement with normal donor-to-donor variability, showing significant interaction with the different treatments (p<0.05). Of note, the YAP and TAZ N/C ratio data acquired on ECM hydrogels closely mirrored results obtained using HTM cells on conventional glass coverslips that served as an additional control (**Suppl. Fig. 3**).

**Fig. 1.**
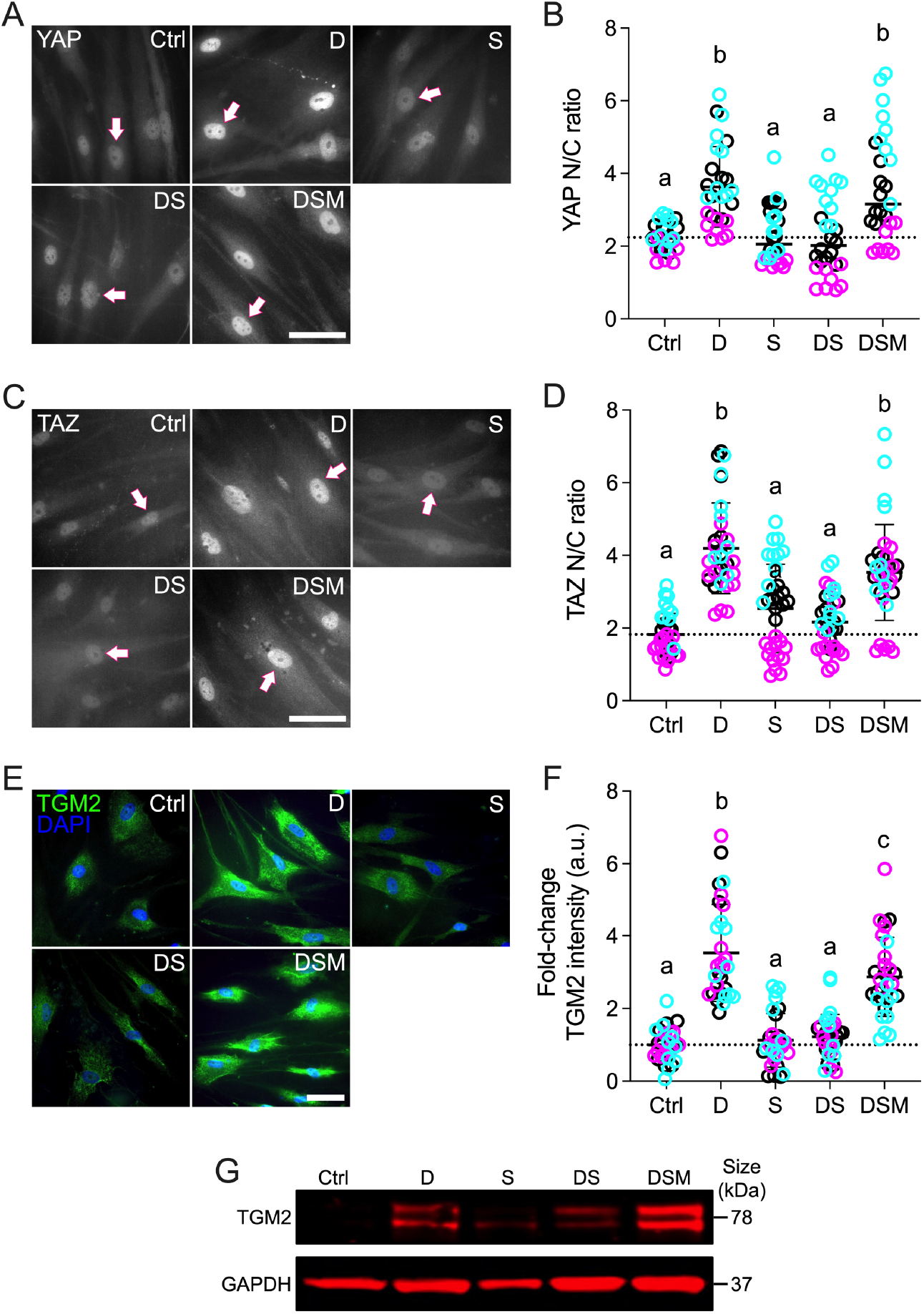
Effects of simvastatin on YAP/TAZ nuclear localization and TGM2 levels in HTM cells. (**A,C,E**) Representative fluorescence micrographs of YAP, TAZ, and TGM2 in HTM cells cultured atop ECM hydrogels subjected to vehicle control, dexamethasone (D; 100 nM), simvastatin (S; 10 μM), dexamethasone + simvastatin, and dexamethasone + simvastatin + mevalonate-5-phosphate (M; 500 μM) at 3 d. Arrows indicate YAP/TAZ nuclear localization. Scale bar, 20 μm. (**B,D,F**) Analysis of YAP/TAZ nuclear/cytoplasmic ratios and TGM2 fluorescence intensity (N = 27-34 images from 3 HTM cell strains with 3 experimental replicates per cell strain). Symbols with different colors represent different cell strains; dotted lines indicate control baselines. The bars and error bars indicate Mean ± SD. Significance was determined by two-way ANOVA using multiple comparisons tests; shared significance indicator letters = non-significant difference (p>0.05), distinct letters = significant difference (p<0.05). (**G**) Qualitative immunoblot of TGM2 with GAPDH serving as loading control (N = 1 per group [pooled from 3 experimental replicates] from 1 HTM cell strain).

Next, we focused on the ECM crosslinking enzyme transglutaminase 2 (TGM2), whose expression is elevated in the TM of glaucomatous eyes^56^. After YAP/TAZ translocate to the nucleus, they interact with TEAD transcription factors to regulate the expression of glaucoma-related putative downstream effectors including TGM2. A previous report showed the inhibition of TGM2 expression in HTM cells with siRNA-mediated YAP knockdown^57^; this observation was recently confirmed in our own study^15^. Consistent with the YAP/TAZ nuclear localization data, dexamethasone treatment significantly increased TGM2 intensity in HTM cells (~3.5-fold) compared to controls (**Fig. 1E,F**); simvastatin alone had no effect on TGM2. Co-treatment with dexamethasone + simvastatin significantly decreased TGM2 intensity (~1.2-fold) compared to dexamethasone-induced HTM cells, restoring baseline levels. The addition of mevalonate-5-phosphate resulted in significantly increased TGM2 intensity (~2.9-fold) compared to dexamethasone + simvastatin, showing comparable levels to the dexamethasone-induced group. While different HTM cell strains did not affect the results (p=0.3848), there was significant interaction with the different treatments (p=0.0070). Qualitative immunoblot analyses validated the TGM2 immunocytochemistry results showing overall very similar trends (**Fig. 1G**).

Taken together, these data demonstrate that simvastatin prevents glucocorticoid-induced pathologic YAP/TAZ nuclear localization and concurrently reduces expression of the downstream effector TGM2 in HTM cells in a tissue-mimetic ECM microenvironment.

### Simvastatin reduces pathologic F-actin and αSMA stress fibers in HTM cells

The actin cytoskeleton is the principal force-generating machinery in the cell^58^. F-actin filaments are sensitive to mechanical stimuli^59^ and have been demonstrated to be essential for YAP/TAZ activity^11,60,61^. In addition, the mevalonate pathway was recently reported to regulate YAP/TAZ through F-actin^62^; therefore, we hypothesized that simvastatin decreases glucocorticoid-induced pathologic actin stress fiber formation in HTM cells. Dexamethasone treatment significantly increased F-actin stress fibers/overall intensity in HTM cells (~2.4-fold) compared to controls; simvastatin alone did not affect F-actin (**Fig. 2A,B**). Co-treatment with dexamethasone + simvastatin significantly decreased F-actin intensity (~0.8-fold) compared to dexamethasone-treated HTM cells, reaching control levels. Supplementation of mevalonate-5-phosphate significantly increased F-actin intensity (~2.0-fold) compared to dexamethasone + simvastatin, equivalent to dexamethasone-induced HTM cells. Different HTM cell strains significantly affected the results (p<0.0001), showing significant interaction with the different treatments (p<0.0001).

**Fig. 2.**
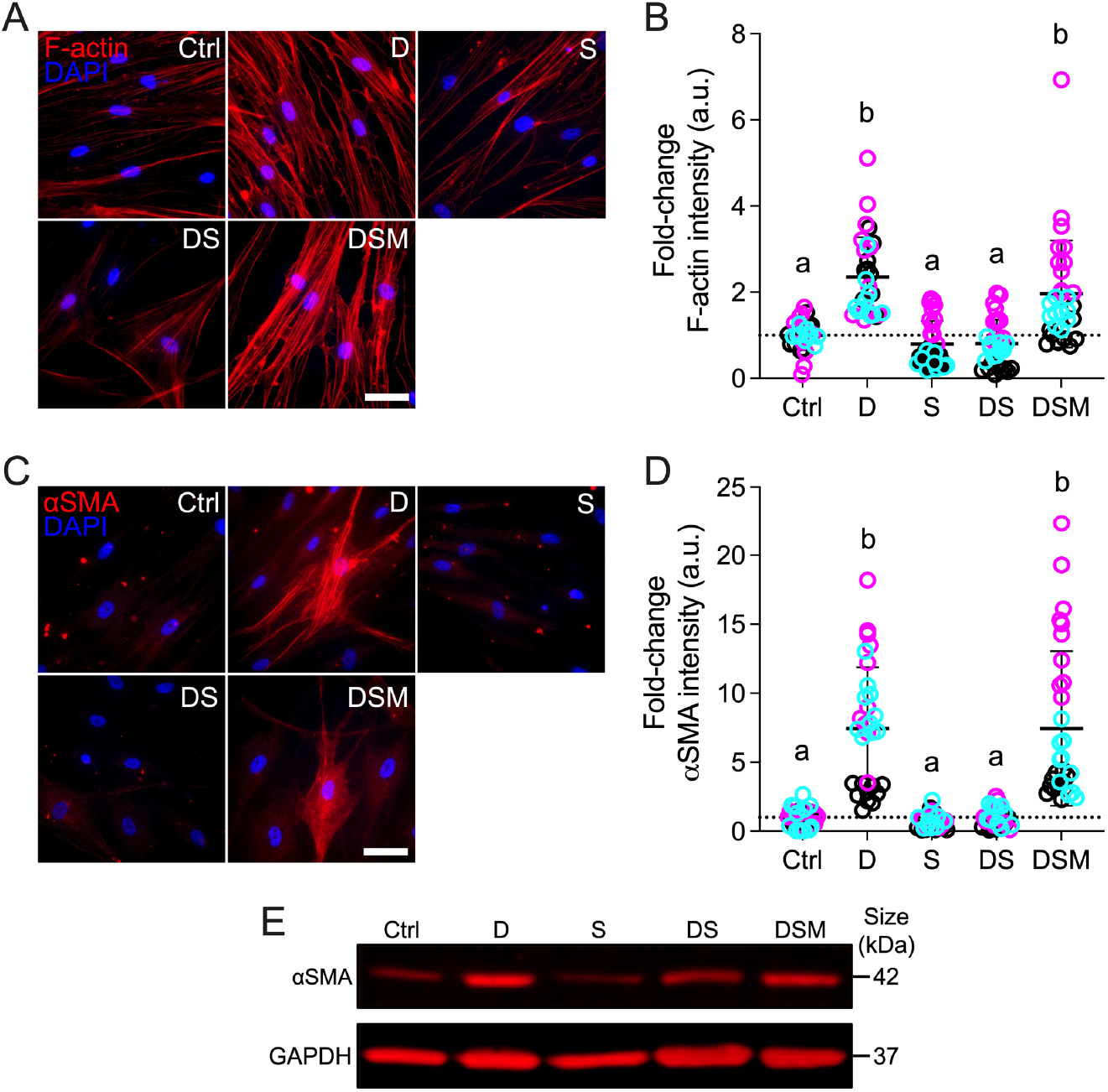
Effects of simvastatin on F-actin and αSMA levels in HTM cells. (**A,C**) Representative fluorescence micrographs of F-actin and αSMA in HTM cells cultured atop ECM hydrogels subjected to vehicle control, dexamethasone (D; 100 nM), simvastatin (S; 10 μM), dexamethasone + simvastatin, and dexamethasone + simvastatin + mevalonate-5-phosphate (M; 500 μM) at 3 d. Scale bar, 20 μm. (**B,D**) Analysis of F-actin and αSMA fluorescence intensities (N = 30-34 images from 3 HTM cell strains with 3 experimental replicates per cell strain). Symbols with different colors represent different cell strains; dotted lines indicate control baselines. The bars and error bars indicate Mean ± SD. Significance was determined by two-way ANOVA using multiple comparisons tests; shared significance indicator letters = non-significant difference (p>0.05), distinct letters = significant difference (p<0.05). (**E**) Qualitative immunoblot of αSMA with GAPDH serving as loading control (N = 1 per group [pooled from 3 experimental replicates] from 1 HTM cell strain).

Next, we looked at alpha-smooth muscle actin (αSMA) - a hallmark of tissue fibrosis^63^; increased αSMA stress fibers have also been linked to fibrotic-like HTM cell pathobiology and outflow dysfunction in glaucoma^64–67^ Consistent with the F-actin data, exposure of HTM cells to dexamethasone significantly increased αSMA stress fibers/overall intensity (~7.5-fold) compared to controls, which only showed very weak staining (**Fig. 2C,D**). Simvastatin alone had no effect on αSMA. Co-treatment with dexamethasone + simvastatin significantly decreased αSMA intensity (~1.0-fold) compared to dexamethasone-induced HTM cells, restoring baseline levels. The addition of mevalonate-5-phosphate significantly increased αSMA stress fibers (~7.5-fold) compared to dexamethasone + simvastatin, identical to dexamethasone-treated HTM cells.

Different HTM cell strains significantly affected the results (p<0.0001), showing significant interaction with the different treatments (p<0.0001). Qualitative immunoblot analyses validated the αSMA immunocytochemistry results showing overall very similar trends (**Fig. 2E**).

Taken together, these data demonstrate that simvastatin blocks glucocorticoid-induced pathologic actin stress fiber formation in HTM cells in a tissue-mimetic ECM microenvironment.

### Simvastatin reduces pathologic p-MLC levels in HTM cells

Myosin light chain is a master regulator of cell contractility when phosphorylated by Rho kinases via the generation of pulling forces from actomyosin filament contraction^68–70^. Increased phospho-myosin light chain (p-MLC) is strongly associated with a pathologic hypercontractile TM cell phenotype akin to activated myofibroblasts^4,47,67^. In contrast, a decrease in p-MLC has been shown to increase aqueous outflow facility in perfusion studies^71^. Statin-mediated inhibition of the mevalonate pathway directly affects downstream Rho GTPase signaling^40^; therefore, we hypothesized that simvastatin decreases glucocorticoid-induced pathologic p-MLC levels in HTM cells. Dexamethasone treatment significantly increased p-MLC intensity in HTM cells (~4.2-fold) compared to controls; simvastatin alone had no effect on p-MLC levels (**Fig. 3A,B**). Co-treatment with dexamethasone + simvastatin significantly decreased p-MLC intensity (~0.8-fold) compared to dexamethasone-induced HTM cells, restoring control levels. The addition of mevalonate-5-phosphate significantly increased p-MLC intensity (~4.3-fold) compared to dexamethasone + simvastatin, indistinguishable from dexamethasone-treated HTM cells. Different HTM cell strains significantly affected the results (p<0.0001), showing significant interaction with the different treatments (p<0.0001).

**Fig. 3.**
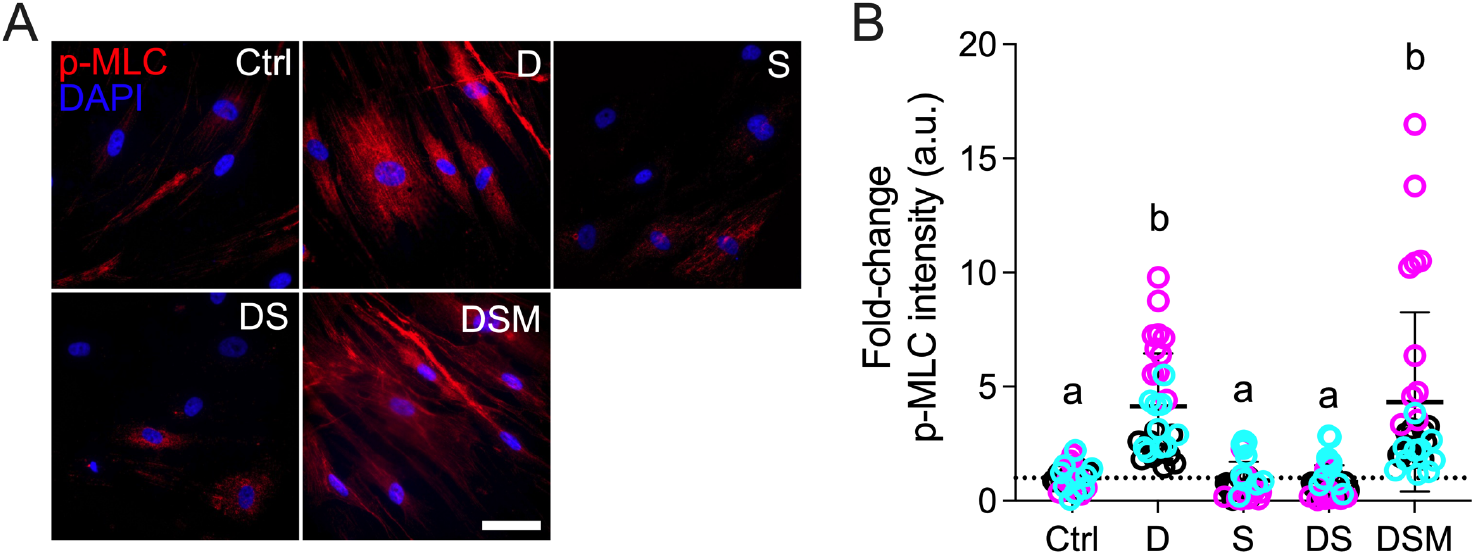
Effect of simvastatin on pMLC levels in HTM cells. (**A**) Representative fluorescence micrographs of p-MLC in HTM cells cultured atop ECM hydrogels subjected to vehicle control, dexamethasone (D; 100 nM), simvastatin (S; 10 μM), dexamethasone + simvastatin, and dexamethasone + simvastatin + mevalonate-5-phosphate (M; 500 μM) at 3 d. Scale bar, 20 μm. (**B**) Analysis of p-MLC fluorescence intensity (N = 29-33 images from 3 HTM cell strains with 3 experimental replicates per cell strain). Symbols with different colors represent different cell strains; dotted line indicates control baseline. The bars and error bars indicate Mean ± SD. Significance was determined by two-way ANOVA using multiple comparisons test; shared significance indicator letters = non-significant difference (p>0.05), distinct letters = significant difference (p<0.05).

From this experiment, we conclude that simvastatin prevents glucocorticoid-induced pathologic myosin light chain phosphorylation in HTM cells in a tissue-mimetic ECM microenvironment to prevent cells from acquiring a hypercontractile myofibroblast-like phenotype.

### Simvastatin reduces pathologic contraction, stiffening, and FN deposition in HTM cells

The TM undergoes increased fibrotic-like contraction and aberrant ECM deposition that together contribute to pathologic tissue stiffening in primary open-angle glaucoma^8^. To better approximate the 3D tissue architecture in the juxtacanalicular TM region that is critical for outflow regulation^9^, HTM cells were encapsulated in ECM hydrogels for tissue-level functional studies. We hypothesized that simvastatin decreases glucocorticoid-induced pathologic contraction, stiffening, and fibronectin deposition in HTM cell-laden hydrogels. Consistent with the p-MLC data and supported by our previous study^46^, exposure of HTM cells to dexamethasone significantly increased hydrogel contraction (~78.1%) compared to controls (**Fig. 4A,B**). Simvastatin alone had no effect on HTM hydrogel contraction. Co-treatment with dexamethasone + simvastatin significantly decreased hydrogel contraction (~103.9%) - or in other words relaxed the constructs - compared to dexamethasone-induced HTM hydrogels, restoring baseline levels.

**Fig. 4.**
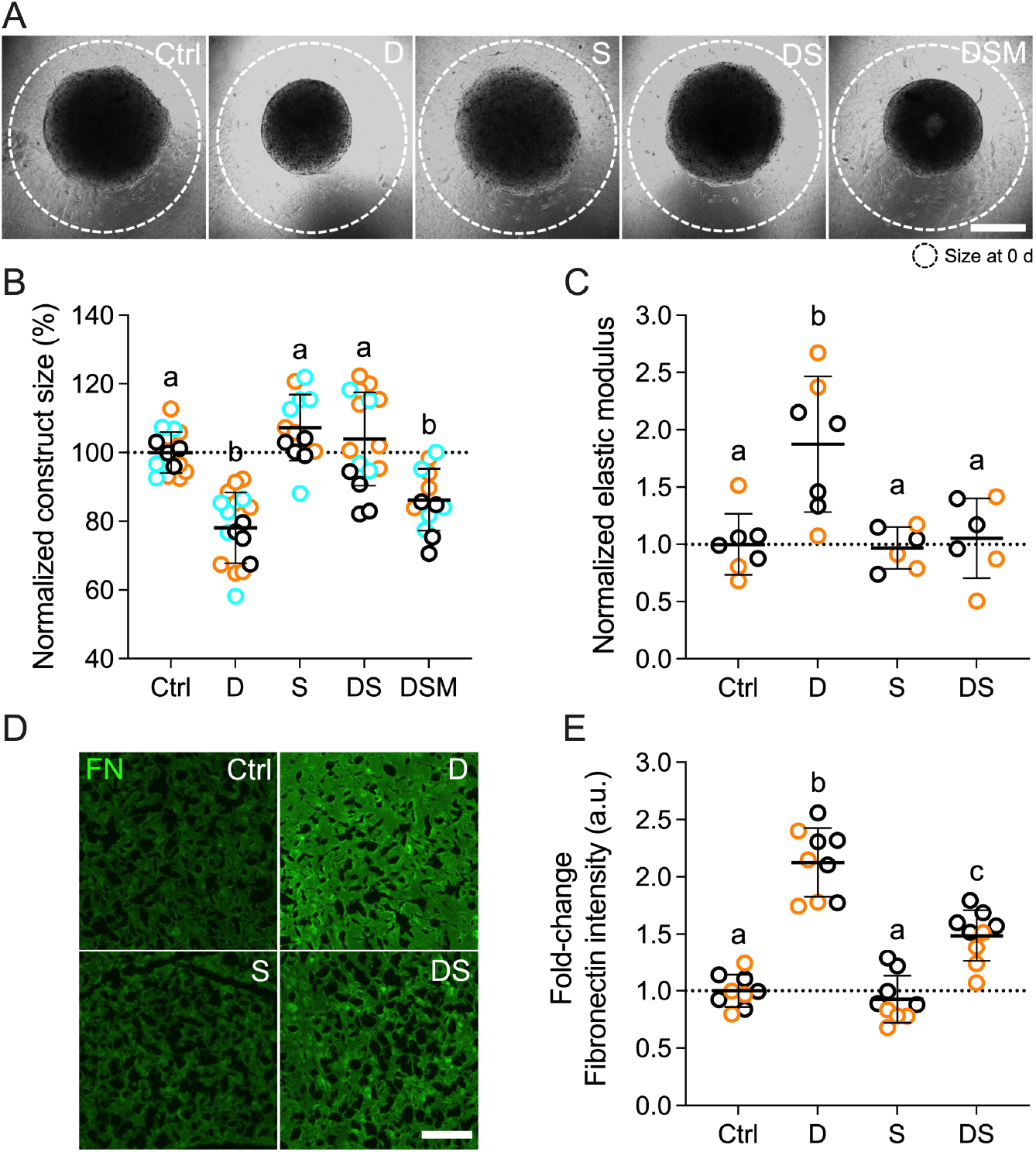
Effects of simvastatin on HTM hydrogel contraction, stiffness, and fibronectin deposition. (**A**) Representative brightfield images of HTM cell-encapsulated ECM hydrogels subjected to vehicle control, dexamethasone (D; 100 nM), simvastatin (S; 10 μM), dexamethasone + simvastatin, and dexamethasone + simvastatin + mevalonate-5-phosphate (M; 500 μM) at 10 d. Dashed lines outline original size of constructs at 0 d. Scale bar, 1 mm. (**B**) Analysis of HTM hydrogel construct size (N = 13-17 experimental replicates from 3 HTM cell strains). (**C**) Analysis of HTM hydrogel elastic modulus (N = 6-7 experimental replicates from 2 HTM cell strains). (**D**) Representative fluorescence micrographs of FN in HTM cell-encapsulated ECM hydrogels. Scale bar, 20 μm. (**E**) Analysis of FN fluorescence intensity (N = 9 images from 2 HTM cell strains with 2 experimental replicates per cell strain). Symbols with different colors represent different cell strains; dotted lines indicate control baselines. The bars and error bars indicate Mean ± SD. Significance was determined by two-way ANOVA using multiple comparisons tests; shared significance indicator letters = non-significant difference (p>0.05), distinct letters = significant difference (p<0.05).

Supplementation of mevalonate-5-phosphate significantly increased hydrogel contraction (~86.2%) compared to dexamethasone + simvastatin, identical to dexamethasone-treated HTM hydrogels. Different HTM cell strains significantly affected the results (p<0.005); yet, there was no significant interaction with the different treatments (p=0.2411). To rule out that hydrogel contractility was influenced by the cell number, HTM cell viability inside the 3D ECM network was assessed. We observed no differences between the different groups (**Suppl. Fig. 4**).

Having shown that the supplementation of mevalonate-5-phosphate fully offsets the inhibitory effects of simvastatin using a variety of techniques – thereby unequivocally confirming that the results seen are mediated by the mevalonate pathway, we dropped the group for subsequent experiments. To assess the functional consequences of increased hydrogel contraction on tissue-level construct stiffness, we next performed oscillatory rheology analyses. Consistent with our previous report^46^, dexamethasone treatment induced significant hydrogel stiffening (~1.9-fold) compared to controls; simvastatin alone had no effect on hydrogel stiffness (**Fig. 4C**). Co-treatment with dexamethasone + simvastatin significantly decreased hydrogel stiffness (~1.1-fold) - or softened the constructs - compared to dexamethasone-induced HTM hydrogels, restoring control levels. Different HTM cell strains did not affect the results (p=0.9700), and there was no significant interaction with the different treatments (p=0.7135).

Lastly, we looked at the deposition of fibronectin (FN) in HTM hydrogels. FN is a major ECM player and signaling component in the native tissue^72^, and has been detected at elevated levels in the TM of glaucomatous eyes^73^. Furthermore, it has long been known that HTM cells express high FN levels upon glaucomatous stimulation^74,75^. Exposure to dexamethasone significantly increased FN intensity (~2.1-fold) compared to controls; simvastatin alone did not affect FN levels (**Fig. 4D,E**). Co-treatment with dexamethasone + simvastatin significantly decreased FN intensity (~1.5-fold) compared to dexamethasone-induced HTM hydrogels, with levels remaining significantly higher than controls. Different HTM cell strains significantly affected the results (p<0.01); yet, there was no significant interaction with the different treatments (p=0.3233).

Taken together, these data demonstrate that simvastatin blocks glucocorticoid-induced pathologic ECM contraction and stiffening, and concurrently reduces extracellular FN deposition in HTM cells in a tissue-mimetic ECM microenvironment.

### Simvastatin rescues pathologic contraction in HTM cells

All experiments to delineate the effects of simvastatin on HTM cell pathobiology up to this point were conducted by simulating a “prophylactic treatment approach”, i.e., dexamethasone and simvastatin were co-delivered from the start. Next, to mimic a “therapeutic treatment approach”, HTM cell-encapsulated hydrogels were treated for 5 days with dexamethasone to establish a precontracted baseline, followed by treatment with simvastatin for 5 days with dexamethasone withheld. Using this design, we assessed the rescue potential of simvastatin in direct comparison with the FDA-approved Rho kinase inhibitor netarsudil, shown to increase aqueous outflow via reducing TM contractile tone^76,77^. We hypothesized that simvastatin rescues glucocorticoid-induced pathologic contraction of HTM cell-laden hydrogels to a similar degree as clinically-used netarsudil. Dexamethasone treatment significantly increased hydrogel contraction (~82.1%) compared to controls (**Fig. 5**). Sequential treatment with simvastatin fully rescued dexamethasone-induced HTM hydrogel contraction (~96.4%), restoring baseline levels. For netarsudil, we observed an even more potent rescuing effect (~112.6%), consistent with our previous data^48^, that exceeded both the simvastatin and control groups. While different HTM cell strains did not affect the results (p=0.2469), there was significant interaction with the different treatments (p<0.0001). HTM cell viability analysis demonstrated that hydrogel contractility was not affected by the cell number; we observed no differences between the different groups (**Suppl. Fig. 5**).

**Fig. 5.**
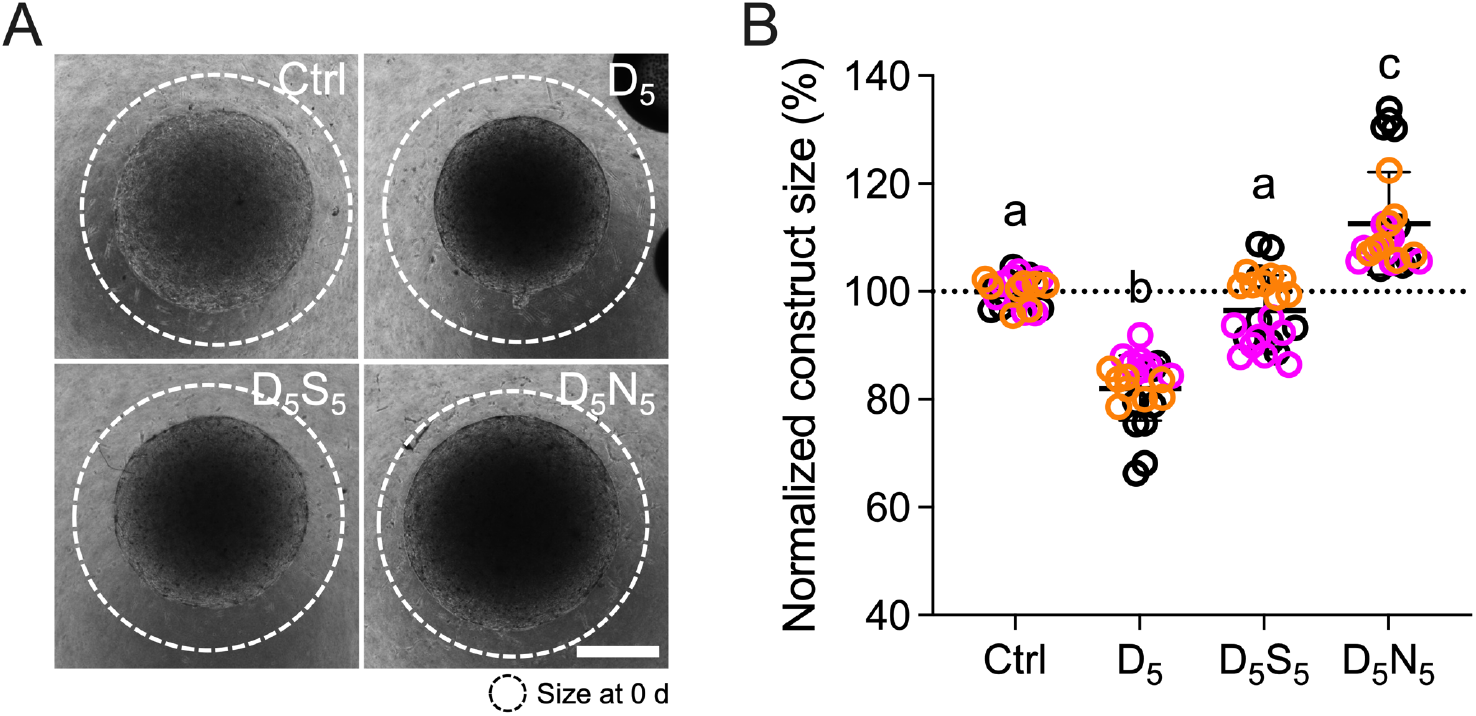
Effect of simvastatin on HTM hydrogel contraction compared to clinically-used Rho kinase inhibitor. (**A**) Representative brightfield images of HTM cell-encapsulated ECM hydrogels subjected to vehicle control, dexamethasone5 (D_5_; 100 nM [0-5 d]; vehicle [5-10 d]), dexamethasone_5_ + simvastatin_5_ (D_5_; [0-5 d]; S_5_; 10 μM [5-10 d]), and dexamethasone_5_ + netarsudil_5_ (D_5_; [0-5 d]; N_5_; 1.0 μM [5-10 d]) at 10 d. Dashed lines outline original size of constructs at 0 d. Scale bar, 1 mm. (**B**) Analysis of HTM hydrogel construct size (N = 24 experimental replicates from 3 HTM cell strains). Symbols with different colors represent different cell strains; dotted line indicates control baseline. The bars and error bars indicate Mean ± SD. Significance was determined by two-way ANOVA using multiple comparisons test; shared significance indicator letters = non-significant difference (p>0.05), distinct letters = significant difference (p<0.05).

From this data we conclude that simvastatin rescues glucocorticoid-induced pathologic ECM contraction in a tissue-mimetic ECM microenvironment, albeit to a lower extent compared to a clinically-used Rho kinase inhibitor.

## Discussion

Simvastatin, a member of the cholesterol-lowering statin drug class, inhibits HMG-CoA reductase that catalyzes the production of mevalonate. The anabolic mevalonate cascade provides key isoprenoid metabolites for diverse cellular processes including cholesterol synthesis and post-translational membrane targeting of Rho GTPases^78,79^. Furthermore, it was recently demonstrated that the mevalonate pathway has a profound impact on the function of the transcriptional regulators YAP and TAZ in different cancer cells^40,62,80^. These studies mechanistically linked the mevalonate pathway to (i) geranylgeranyl pyrophosphate (GGPP)-mediated Rho GTPase activation and F-actin fiber assembly rather than the squalene/cholesterol arm of the mevalonate cascade, and (ii) reduction of YAP/TAZ inhibitory phosphorylation and sustained YAP/TAZ transcriptional activity via nuclear accumulation independent of canonical Hippo-LATS1/2 kinase activity. With the increasing evidence - including from our own laboratory - of aberrant YAP/TAZ activity in human TM cells isolated from glaucoma eyes or induced with glaucoma-associated stressors^15–22^, in conjunction with the recent discovery of YAP as a potential “risk gene” for open-angle glaucoma^23^, we here tested the hypothesis that simvastatin decreases YAP/TAZ activity in glucocorticoid-induced TM cells to attenuate glaucomatous cell pathobiology in a tissue-mimetic ECM microenvironment^46^.

Simvastatin was identified among more than 600 FDA-approved compounds to have potent inhibitory effects on YAP/TAZ activity in human breast cancer cells, and to efficiently rescue *Drosophila* eye overgrowth induced by the YAP orthologue *Yki*^40^. In another high-throughput library screen of more than 13,000 small-molecule compounds, simvastatin was found to exhibit strong YAP inhibitory effects via the same GGPP/Rho/actin signaling axis in human lung fibroblasts, and to reduce fibrotic markers in the bleomycin mouse model of pulmonary fibrosis^81^. Together, these data support the notion that simvastatin has intriguing potential to modulate pathologic YAP/TAZ activity in the context of TM cell dysfunction that has been associated with fibrotic-like ECM remodeling, tissue contraction and stiffening in glaucoma^7,8^. We found that simvastatin potently decreased dexamethasone-induced YAP/TAZ nuclear localization in HTM cells, the main mechanism to regulate their function^11^, concurrent with a reduction of the downstream effector TGM2 known to be expressed at increased levels in the TM of glaucoma eyes^56^. These data were in agreement with our recent study in which we showed that genetic or pharmacologic YAP/TAZ inactivation potently blocked TGFß2-induced HTM cell pathobiology in the same ECM hydrogel environment^15^. Of note, we observed nearly identical trends for YAP and TAZ. This suggests that simvastatin may regulate the two paralogues in a similar manner in HTM cells consistent with their acknowledged functional redundancy^82^. We also found that the simvastatin-mediated YAP/TAZ inactivation in HTM cells was fully negated when mevalonate-5-phosphate was supplemented, comparable to what has been reported in non-ocular cells^40^. While we did not inhibit distinct enzymes here to identify the specific metabolic intermediate involved in YAP/TAZ regulation in HTM cells, evidence from previous studies strongly support that protein geranylgeranylation is responsible for the positive effect of the mevalonate pathway on YAP/TAZ activity^40,62,80,81^. It was shown that only farnesyl diphosphate synthase or geranylgeranyl transferase inhibition were able to reproduce the effect of simvastatin on YAP/TAZ nuclear localization and transcriptional activity, whereas inhibition of squalene synthase - the enzyme that catalyzes the first step of sterol biosynthesis - or farnesyl transferase had no effect^40^.

YAP/TAZ activity requires actomyosin cytoskeletal integrity/tension and involves contractile as well as adhesive structures^83,84^ We found that simvastatin effectively blocked glucocorticoid-induced F-actin stress fiber formation in HTM cells. This was in agreement with our recent study using verteporfin to inhibit YAP/TAZ^15^ and data demonstrating that the mevalonate pathway regulates YAP/TAZ through F-actin^62^. Previously, HTM cells treated with lovastatin or geranylgeranyl transferase inhibitor were shown to exhibit decreased actin stress fiber organization and increased accumulation of unprenylated (i.e., inactive) RhoA and RhoB^35^, lending further support for protein geranylgeranylation as being the nexus of statin-mediated effects on HTM cell cytoskeletal organization. We also observed that simvastatin treatment potently prevented αSMA fiber formation, an accepted indicator of a fibrotic-like HTM cell phenotype^67^. These data were consistent with evidence that simvastatin exhibits anti-fibrotic potential in human lung fibroblasts by targeting the GGPP/Rho/actin signaling axis^81^.

The Rho/ROCK pathway is a master regulator of the actin cytoskeleton that has been strongly associated with HTM cell contractility via phosphorylation of myosin light chain^67^. We showed that simvastatin potently decreased dexamethasone-induced p-MLC levels in a tissue-mimetic ECM environment to prevent HTM cells from acquiring a hypercontractile myofibroblastlike phenotype. Consistent with the p-MLC data, we further demonstrated that simvastatin efficiently blocked pathologic contraction and stiffening of HTM cell-encapsulated ECM hydrogels that more accurately simulate the native tissue architecture. These functional alterations were found to coincide with a reduction in FN deposition that is known to play a role in the development of ocular hypertension/glaucoma^72,73^. These cumulative findings are in line with our previous observations that YAP/TAZ inhibition with verteporfin counteracts tissue-level functional impairments^15^, further strengthening the argument that aberrant YAP/TAZ activity in cells of the outflow tract may contribute to glaucoma pathogenesis. Importantly, our data suggest that supplementation of simvastatin can both prevent (i.e., co-treatment) and rescue (i.e., sequential treatment) dexamethasone-induced TM cell pathobiology contingent on cell-ECM interactions. We acknowledge that simvastatin fell short of matching the potent contraction-reversing effects of the FDA-approved ROCK inhibitor netarsudil, the active ingredient in Rhopressa™, that increases outflow through the stiffened TM via reducing tissue contraction as a function of ECM-focal adhesion and actin stress fiber disassembly^76,77,85–87^. This could be explained by the direct vs. indirect modes of action on cellular contractility. Potential co-treatment regimens could be explored in future studies.

In conclusion, we demonstrated that YAP/TAZ inactivation with simvastatin attenuates dexamethasone-induced HTM cell dysfunction in a tissue-mimetic ECM microenvironment. Our data may help explain the association of statin use with a reduced risk of developing glaucoma by proposing indirect YAP/TAZ inhibition as a regulatory mechanism, and highlight the therapeutic potential of localized simvastatin therapy to treat outflow tissue dysfunction in glaucoma.

## Disclosure

The authors report no conflicts of interest.

## Funding

This project was supported in part by National Institutes of Health grants K08EY031755 (to P.S.G), an American Glaucoma Society Young Clinician Scientist Award (to P.S.G.), a Syracuse University BioInspired Pilot Grant (to S.H.), unrestricted grants to SUNY Upstate Medical University Department of Ophthalmology and Visual Sciences from Research to Prevent Blindness (RPB) and from Lions Region 20-Y1, and RPB Career Development Awards (to P.S.G. and S.H.).

## Acknowledgments

We thank Dr. Robert W. Weisenthal and the team at Specialty Surgery Center of Central New York for assistance with corneal rim specimens. We also thank Dr. Nasim Annabi at the University of California – Los Angeles for providing the KCTS-ELP, Dr. Alison Patteson at Syracuse University for rheometer access, and Drs. Audrey M. Bernstein and Mariano S. Viapiano at Upstate Medical University for imaging support.

## Author contributions

H.Y., A.S., H.L., A.N.S., T.B., P.S.G., and S.H. designed all experiments, collected, analyzed, and interpreted the data. H.Y., A.S., and S.H. wrote the manuscript. All authors commented on and approved the final manuscript. P.S.G. and S.H. conceived and supervised the research.

## Competing interests

The authors declare no conflict of interest.

## Data and materials availability

All data needed to evaluate the conclusions in the paper are present in the paper and/or the Supplementary Materials. Additional data related to this paper may be requested from the authors.

## Notes

### Competing Interest Statement

The authors have declared no competing interest.

## References

1. Brubaker RF. Flow of aqueous humor in humans [The Friedenwald Lecture]. Invest Ophthalmol Vis Sci 1991;32:3145–3166.

2. Tamm ER. The trabecular meshwork outflow pathways: structural and functional aspects. Exp Eye Res 2009;88:648–655.

3. Tamm ER, Braunger BM, Fuchshofer R. Intraocular Pressure and the Mechanisms Involved in Resistance of the Aqueous Humor Flow in the Trabecular Meshwork Outflow Pathways. Prog Mol Biol Transl Sci 2015;134:301–314.

4. Stamer WD, Clark AF. The many faces of the trabecular meshwork cell. Exp Eye Res 2017;158:112–123.

5. Acott TS, Kelley MJ. Extracellular matrix in the trabecular meshwork. Exp Eye Res 2008;86:543–561.

6. Kelley MJ, Rose AY, Keller KE, Hessle H, Samples JR, Acott TS. Stem cells in the trabecular meshwork: present and future promises. Exp Eye Res 2009;88:747–751.

7. Kwon YH, Fingert JH, Kuehn MH, Alward WL. Primary open-angle glaucoma. N Engl J Med 2009;360:1113–1124.

8. Wang K, Read AT, Sulchek T, Ethier CR. Trabecular meshwork stiffness in glaucoma. Exp Eye Res 2017;158:3–12.

9. Stamer WD, Acott TS. Current understanding of conventional outflow dysfunction in glaucoma. Curr Opin Ophthalmol 2012;23:135–143.

10. Acott TS, Vranka JA, Keller KE, Raghunathan V, Kelley MJ. Normal and glaucomatous outflow regulation. Prog Retin Eye Res 2021;82:100897.

11. Dupont S, Morsut L, Aragona M, et al. Role of YAP/TAZ in mechanotransduction. Nature 2011;474:179–183.

12. Piccolo S, Dupont S, Cordenonsi M. The biology of YAP/TAZ: hippo signaling and beyond. Physiol Rev 2014;94:1287–1312.

13. Varelas X. The Hippo pathway effectors TAZ and YAP in development, homeostasis and disease. Development 2014;141:1614–1626.

14. Panciera T, Azzolin L, Cordenonsi M, Piccolo S. Mechanobiology of YAP and TAZ in physiology and disease. Nat Rev Mol Cell Biol 2017;18:758–770.

15. Li H, Raghunathan V, Stamer WD, Ganapathy PS, Herberg S. Extracellular Matrix Stiffness and TGFbeta2 Regulate YAP/TAZ Activity in Human Trabecular Meshwork Cells. Front Cell Dev Biol 2022;10:844342.

16. Yemanyi F, Vranka J, Raghunathan VK. Crosslinked Extracellular Matrix Stiffens Human Trabecular Meshwork Cells Via Dysregulating beta-catenin and YAP/TAZ Signaling Pathways. Invest Ophthalmol Vis Sci 2020;61:41.

17. Peng J, Wang H, Wang X, Sun M, Deng S, Wang Y. YAP and TAZ mediate steroid-induced alterations in the trabecular meshwork cytoskeleton in human trabecular meshwork cells. Int J Mol Med 2018;41:164–172.

18. Chen WS, Cao Z, Krishnan C, Panjwani N. Verteporfin without light stimulation inhibits YAP activation in trabecular meshwork cells: Implications for glaucoma treatment. Biochem Biophys Res Commun 2015;466:221–225.

19. Ho LTY, Skiba N, Ullmer C, Rao PV. Lysophosphatidic Acid Induces ECM Production via Activation of the Mechanosensitive YAP/TAZ Transcriptional Pathway in Trabecular Meshwork Cells. Invest Ophthalmol Vis Sci 2018;59:1969–1984.

20. Dhamodaran K, Baidouri H, Sandoval L, Raghunathan V. Wnt Activation After Inhibition Restores Trabecular Meshwork Cells Toward a Normal Phenotype. Invest Ophthalmol Vis Sci 2020;61:30.

21. Yemanyi F, Raghunathan V. Lysophosphatidic Acid and IL-6 Trans-signaling Interact via YAP/TAZ and STAT3 Signaling Pathways in Human Trabecular Meshwork Cells. Invest Ophthalmol Vis Sci 2020;61:29.

22. Thomasy SM, Morgan JT, Wood JA, Murphy CJ, Russell P. Substratum stiffness and latrunculin B modulate the gene expression of the mechanotransducers YAP and TAZ in human trabecular meshwork cells. Exp Eye Res 2013;113:66–73.

23. Gharahkhani P, Jorgenson E, Hysi P, et al. Genome-wide meta-analysis identifies 127 open-angle glaucoma loci with consistent effect across ancestries. Nat Commun 2021;12:1258.

24. Sirtori CR. The pharmacology of statins. Pharmacol Res 2014;88:3–11.

25. DeBose-Boyd RA. Feedback regulation of cholesterol synthesis: sterol-accelerated ubiquitination and degradation of HMG CoA reductase. Cell Res 2008;18:609–621.

26. Posch-Pertl L, Michelitsch M, Wagner G, et al. Cholesterol and glaucoma: a systematic review and meta-analysis. Acta Ophthalmol 2022;100:148–158.

27. Wang S, Bao X. Hyperlipidemia, Blood Lipid Level, and the Risk of Glaucoma: A Meta-Analysis. Invest Ophthalmol Vis Sci 2019;60:1028–1043.

28. McGwin G, Jr., McNeal S, Owsley C, Girkin C, Epstein D, Lee PP. Statins and other cholesterol-lowering medications and the presence of glaucoma. Arch Ophthalmol 2004;122:822–826.

29. Talwar N, Musch DC, Stein JD. Association of Daily Dosage and Type of Statin Agent With Risk of Open-Angle Glaucoma. JAMA Ophthalmol 2017;135:263–267.

30. De Castro DK, Punjabi OS, Bostrom AG, et al. Effect of statin drugs and aspirin on progression in open-angle glaucoma suspects using confocal scanning laser ophthalmoscopy. Clin Exp Ophthalmol 2007;35:506–513.

31. McCann P, Hogg RE, Fallis R, Azuara-Blanco A. The Effect of Statins on Intraocular Pressure and on the Incidence and Progression of Glaucoma: A Systematic Review and Meta-Analysis. Invest Ophthalmol Vis Sci 2016;57:2729–2748.

32. Stein JD, Newman-Casey PA, Talwar N, Nan B, Richards JE, Musch DC. The relationship between statin use and open-angle glaucoma. Ophthalmology 2012;119:2074–2081.

33. Thiermeier N, Lammer R, Mardin C, Hohberger B. Erlanger Glaucoma Registry: Effect of a Long-Term Therapy with Statins and Acetyl Salicylic Acid on Glaucoma Conversion and Progression. Biology (Basel) 2021;10.

34. Song J, Deng PF, Stinnett SS, Epstein DL, Rao PV. Effects of cholesterol-lowering statins on the aqueous humor outflow pathway. Invest Ophthalmol Vis Sci 2005;46:2424–2432.

35. Von Zee CL, Richards MP, Bu P, Perlman JI, Stubbs EB, Jr. Increased RhoA and RhoB protein accumulation in cultured human trabecular meshwork cells by lovastatin. Invest Ophthalmol Vis Sci 2009;50:2816–2823.

36. Von Zee CL, Stubbs EB, Jr. Geranylgeranylation facilitates proteasomal degradation of rho G-proteins in human trabecular meshwork cells. Invest Ophthalmol Vis Sci 2011;52:1676–1683.

37. Rodriguez F, Maron DJ, Knowles JW, Virani SS, Lin S, Heidenreich PA. Association Between Intensity of Statin Therapy and Mortality in Patients With Atherosclerotic Cardiovascular Disease. JAMA Cardiol 2017;2:47–54.

38. Song XY, Chen YY, Liu WT, et al. Atorvastatin reduces IOP in ocular hypertension in vivo and suppresses ECM in trabecular meshwork perhaps via FGD4. Int J Mol Med 2022;49.

39. Cong L, Fu S, Zhang J, Zhao J, Zhang Y. Effects of atorvastatin on porcine aqueous humour outflow and trabecular meshwork cells. Exp Ther Med 2018;15:210–216.

40. Sorrentino G, Ruggeri N, Specchia V, et al. Metabolic control of YAP and TAZ by the mevalonate pathway. Nat Cell Biol 2014;16:357–366.

41. Oku Y, Nishiya N, Shito T, et al. Small molecules inhibiting the nuclear localization of YAP/TAZ for chemotherapeutics and chemosensitizers against breast cancers. FEBS Open Bio 2015;5:542–549.

42. Wang K, Li G, Read AT, et al. The relationship between outflow resistance and trabecular meshwork stiffness in mice. Sci Rep 2018;8:5848.

43. Jones R, 3rd, Rhee DJ. Corticosteroid-induced ocular hypertension and glaucoma: a brief review and update of the literature. Curr Opin Ophthalmol 2006;17:163–167.

44. Raghunathan VK, Morgan JT, Park SA, et al. Dexamethasone Stiffens Trabecular Meshwork, Trabecular Meshwork Cells, and Matrix. Invest Ophthalmol Vis Sci 2015;56:4447–4459.

45. Clark AF, Wilson K, McCartney MD, Miggans ST, Kunkle M, Howe W. Glucocorticoid-induced formation of cross-linked actin networks in cultured human trabecular meshwork cells. Invest Ophthalmol Vis Sci 1994;35:281–294.

46. Li H, Bague T, Kirschner A, et al. A tissue-engineered human trabecular meshwork hydrogel for advanced glaucoma disease modeling. Exp Eye Res 2021;205:108472.

47. Li H, Henty-Ridilla JL, Bernstein AM, Ganapathy PS, Herberg S. TGFbeta2 regulates human trabecular meshwork cell contractility via ERK and ROCK pathways with distinct signaling crosstalk dependent on the culture substrate. Curr Eye Res 2022;1–41.

48. Bagué T, Singh A, Ghosh R, et al. Effects of Netarsudil-Family Rho Kinase Inhibitors on Human Trabecular Meshwork Cell Contractility and Actin Remodeling Using a Bioengineered ECM Hydrogel. Frontiers in Ophthalmology 2022;2.

49. Stamer WD, Seftor RE, Williams SK, Samaha HA, Snyder RW. Isolation and culture of human trabecular meshwork cells by extracellular matrix digestion. Curr Eye Res 1995;14:611–617.

50. Keller KE, Bhattacharya SK, Borras T, et al. Consensus recommendations for trabecular meshwork cell isolation, characterization and culture. Exp Eye Res 2018;171:164–173.

51. Zhang YN, Avery RK, Vallmajo-Martin Q, et al. A Highly Elastic and Rapidly Crosslinkable Elastin-Like Polypeptide-Based Hydrogel for Biomedical Applications. Adv Funct Mater 2015;25:4814–4826.

52. Schindelin J, Arganda-Carreras I, Frise E, et al. Fiji: an open-source platform for biological-image analysis. Nat Methods 2012;9:676–682.

53. Timothy P. Lodge PCH. Polymer Chemistry. CRC Press 2020.

54. Kwon H, Kim J, Jho EH. Role of the Hippo pathway and mechanisms for controlling cellular localization of YAP/TAZ. FEBS J 2021.

55. Li H, Perkumas KM, Stamer WD, Ganapathy PS, Herberg S. YAP/TAZ mediate TGFβ2-induced Schlemm’s canal cell dysfunction. bioRxiv 2022;2022.2006.2006.494681.

56. Tovar-Vidales T, Roque R, Clark AF, Wordinger RJ. Tissue transglutaminase expression and activity in normal and glaucomatous human trabecular meshwork cells and tissues. Invest Ophthalmol Vis Sci 2008;49:622–628.

57. Raghunathan VK, Morgan JT, Dreier B, et al. Role of substratum stiffness in modulating genes associated with extracellular matrix and mechanotransducers YAP and TAZ. Invest Ophthalmol Vis Sci 2013;54:378–386.

58. Svitkina T. The Actin Cytoskeleton and Actin-Based Motility. Cold Spring Harb Perspect Biol 2018;10.

59. Dasgupta I, McCollum D. Control of cellular responses to mechanical cues through YAP/TAZ regulation. J Biol Chem 2019;294:17693–17706.

60. Wada K, Itoga K, Okano T, Yonemura S, Sasaki H. Hippo pathway regulation by cell morphology and stress fibers. Development 2011;138:3907–3914.

61. Mo JS, Yu FX, Gong R, Brown JH, Guan KL. Regulation of the Hippo-YAP pathway by protease-activated receptors (PARs). Genes Dev 2012;26:2138–2143.

62. Wang Z, Wu Y, Wang H, et al. Interplay of mevalonate and Hippo pathways regulates RHAMM transcription via YAP to modulate breast cancer cell motility. Proc Natl Acad Sci U S A 2014;111:E89–98.

63. Hinz B, Celetta G, Tomasek JJ, Gabbiani G, Chaponnier C. Alpha-smooth muscle actin expression upregulates fibroblast contractile activity. Mol Biol Cell 2001;12:2730–2741.

64. Tamm ER, Siegner A, Baur A, Lutjen-Drecoll E. Transforming growth factor-beta 1 induces alpha-smooth muscle-actin expression in cultured human and monkey trabecular meshwork. Exp Eye Res 1996;62:389–397.

65. Li G, Lee C, Read AT, et al. Anti-fibrotic activity of a rho-kinase inhibitor restores outflow function and intraocular pressure homeostasis. Elife 2021;10.

66. Honjo M, Igarashi N, Nishida J, et al. Role of the Autotaxin-LPA Pathway in Dexamethasone-Induced Fibrotic Responses and Extracellular Matrix Production in Human Trabecular Meshwork Cells. Invest Ophthalmol Vis Sci 2018;59:21–30.

67. Pattabiraman PP, Rao PV. Mechanistic basis of Rho GTPase-induced extracellular matrix synthesis in trabecular meshwork cells. Am J Physiol Cell Physiol 2010;298:C749–763.

68. Amano M, Nakayama M, Kaibuchi K. Rho-kinase/ROCK: A key regulator of the cytoskeleton and cell polarity. Cytoskeleton (Hoboken) 2010;67:545–554.

69. Wang J, Liu X, Zhong Y. Rho/Rho-associated kinase pathway in glaucoma (Review). Int J Oncol 2013;43:1357–1367.

70. Lampi MC, Reinhart-King CA. Targeting extracellular matrix stiffness to attenuate disease: From molecular mechanisms to clinical trials. Sci Transl Med 2018;10.

71. Rao PV, Deng P, Sasaki Y, Epstein DL. Regulation of myosin light chain phosphorylation in the trabecular meshwork: role in aqueous humour outflow facility. Exp Eye Res 2005;80:197–206.

72. Faralli JA, Schwinn MK, Gonzalez JM, Jr., Filla MS, Peters DM. Functional properties of fibronectin in the trabecular meshwork. Exp Eye Res 2009;88:689–693.

73. Faralli JA, Filla MS, Peters DM. Role of Fibronectin in Primary Open Angle Glaucoma. Cells 2019;8.

74. Zhou L, Li Y, Yue BY. Glucocorticoid effects on extracellular matrix proteins and integrins in bovine trabecular meshwork cells in relation to glaucoma. Int J Mol Med 1998;1:339–346.

75. Medina-Ortiz WE, Belmares R, Neubauer S, Wordinger RJ, Clark AF. Cellular fibronectin expression in human trabecular meshwork and induction by transforming growth factor-beta2. Invest Ophthalmol Vis Sci 2013;54:6779–6788.

76. Tanna AP, Johnson M. Rho Kinase Inhibitors as a Novel Treatment for Glaucoma and Ocular Hypertension. Ophthalmology 2018;125:1741–1756.

77. Rao PV, Pattabiraman PP, Kopczynski C. Role of the Rho GTPase/Rho kinase signaling pathway in pathogenesis and treatment of glaucoma: Bench to bedside research. Exp Eye Res 2017;158:23–32.

78. Goldstein JL, Brown MS. Regulation of the mevalonate pathway. Nature 1990;343:425–430.

79. Casey PJ, Seabra MC. Protein prenyltransferases. J Biol Chem 1996;271:5289–5292.

80. Tanaka K, Osada H, Murakami-Tonami Y, Horio Y, Hida T, Sekido Y. Statin suppresses Hippo pathway-inactivated malignant mesothelioma cells and blocks the YAP/CD44 growth stimulatory axis. Cancer Lett 2017;385:215–224.

81. Santos DM, Pantano L, Pronzati G, et al. Screening for YAP Inhibitors Identifies Statins as Modulators of Fibrosis. Am J Respir Cell Mol Biol 2020;62:479–492.

82. Totaro A, Panciera T, Piccolo S. YAP/TAZ upstream signals and downstream responses. Nature Cell Biology 2018;20:888–899.

83. Valon L, Marin-Llaurado A, Wyatt T, Charras G, Trepat X. Optogenetic control of cellular forces and mechanotransduction. Nat Commun 2017;8:14396.

84. Das A, Fischer RS, Pan D, Waterman CM. YAP Nuclear Localization in the Absence of Cell-Cell Contact Is Mediated by a Filamentous Actin-dependent, Myosin II- and Phospho-YAP-independent Pathway during Extracellular Matrix Mechanosensing. J Biol Chem 2016;291:6096–6110.

85. Rao PV, Deng PF, Kumar J, Epstein DL. Modulation of aqueous humor outflow facility by the Rho kinase-specific inhibitor Y-27632. Invest Ophthalmol Vis Sci 2001;42:1029–1037.

86. Wang SK, Chang RT. An emerging treatment option for glaucoma: Rho kinase inhibitors. Clin Ophthalmol 2014;8:883–890.

87. Zhang K, Zhang L, Weinreb RN. Ophthalmic drug discovery: novel targets and mechanisms for retinal diseases and glaucoma. Nat Rev Drug Discov 2012;11:541–559.

