## Supplemental figures for "YAP/TAZ inactivation with simvastatin attenuates glucocorticoid-induced human trabecular meshwork cell dysfunction"

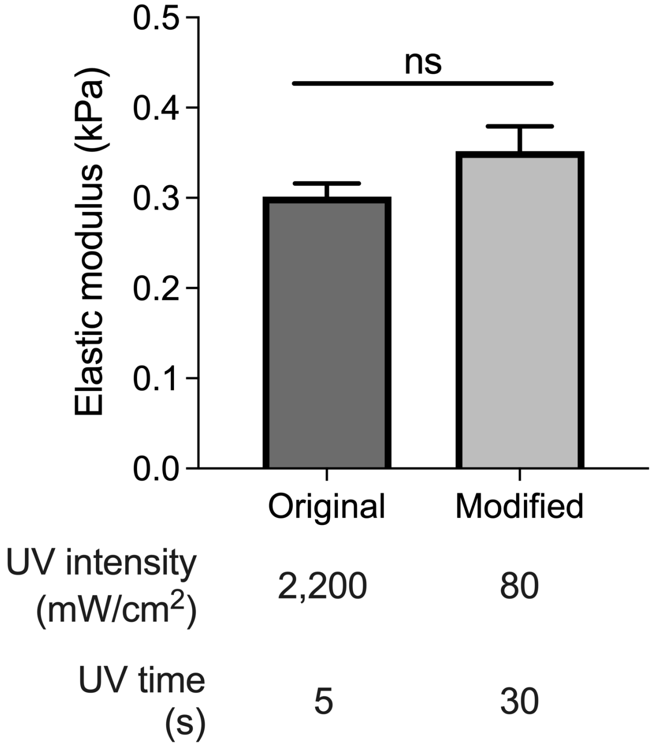


**Suppl. Fig. 1. Effect of UV crosslinking conditions on ECM hydrogel stiffness.** Analysis of acellular ECM hydrogel elastic modulus (N = 3 experimental replicates) as a function of different UV settings. The bars and error bars indicate Mean ± SD. Significance was determined by unpaired *t* test; ns = not significant.

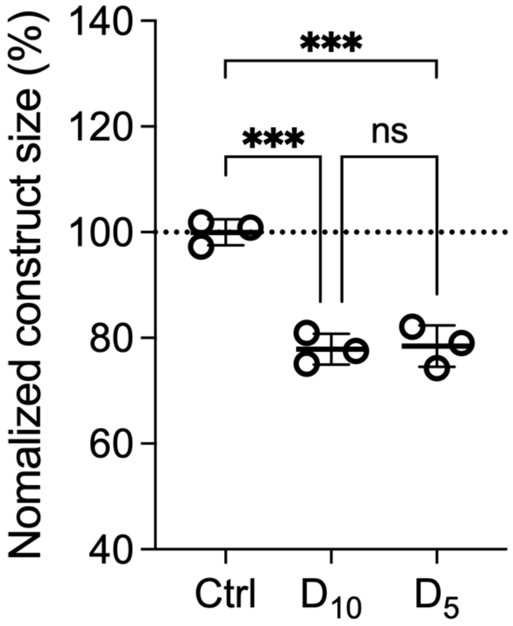


**Suppl. Fig. 2. Effect of dexamethasone treatment on HTM hydrogel contraction.** Analysis of HTM cell-encapsulated ECM hydrogel construct size (N = 3 experimental replicates from 1 HTM cell strain) subjected to vehicle control, dexamethasone$_{10}$ (D; 100 nM [0-10 d]), or dexamethasone$_{5}$ (D; 100 nM [0-5 d]; vehicle [5-10 d]) at 10 d. The bars and error bars indicate Mean ± SD. Significance was determined by one-way ANOVA using multiple comparisons tests; ***$p<0.001$; ns = not significant.

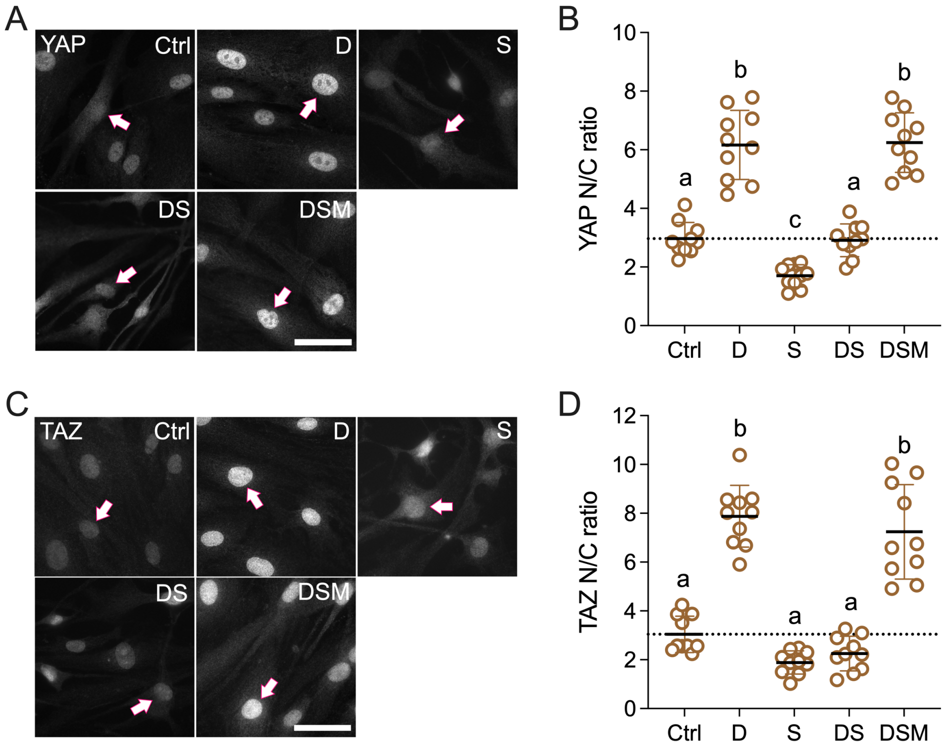


**Suppl. Fig. 3. Effects of simvastatin on YAP/TAZ nuclear localization in HTM cells on glass.** (**A,C**) Representative fluorescence micrographs of YAP and TAZ in HTM cells cultured atop conventional glass coverslips subjected to vehicle control, dexamethasone (D; 100 nM), simvastatin (S; 10 μM), dexamethasone + simvastatin, and dexamethasone + simvastatin + mevalonate-5-phosphate (M; 500 μM) at 3 d. Arrows indicate YAP/TAZ nuclear localization. Scale bar, 20 μm. (**B,D**) Analysis of YAP/TAZ nuclear/cytoplasmic ratios (N = 10 images from 1 HTM cell strain with 3 experimental replicates). Dotted lines indicate control baselines. The bars and error bars indicate Mean ± SD. Significance was determined by one-way ANOVA using multiple comparisons tests; shared significance indicator letters = non-significant difference ($p>0.05$), distinct letters = significant difference ($p<0.05$).

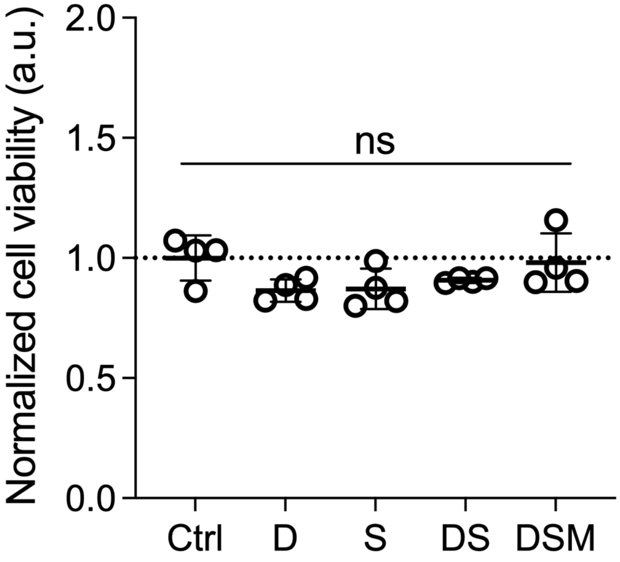


**Suppl. Fig. 4. Effect of different experimental treatments on HTM cell viability.** Analysis of ECM hydrogel-encapsulated HTM cell viability (N = 4 experimental replicates from 1 HTM cell strain) subjected to vehicle control, dexamethasone (D; 100 nM), simvastatin (S; 10 µM), dexamethasone + simvastatin, and dexamethasone + simvastatin + mevalonate-5-phosphate (M; 500 µM) at 10 d. The bars and error bars indicate Mean ± SD. Significance was determined by one-way ANOVA using multiple comparisons test; ns = not significant.

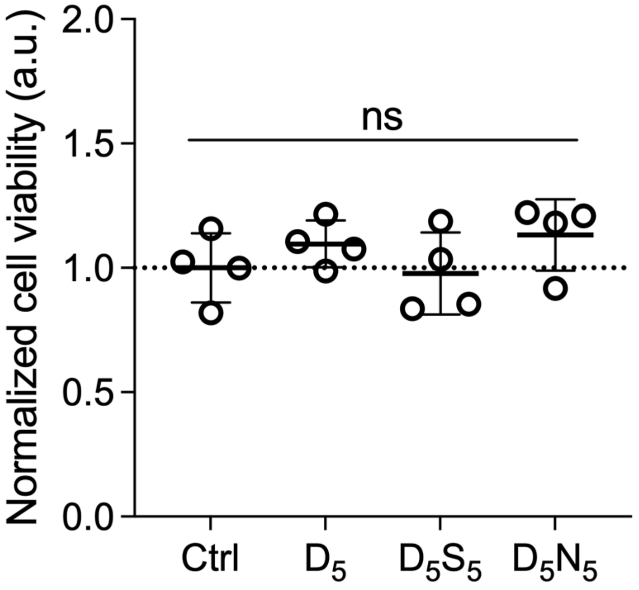


**Suppl. Fig. 5. Effect of different experimental treatments on HTM cell viability.** Analysis of ECM hydrogel-encapsulated HTM cell viability (N = 4 experimental replicates from 1 HTM cell strain) subjected to vehicle control, dexamethasone$_5$ ($D_5$; 100 nM [0-5 d]; vehicle [5-10 d]), dexamethasone$_5$ + simvastatin$_5$ ($D_5$; [0-5 d]; $S_5$; 10 µM [5-10 d]), and dexamethasone$_5$ + netarsudil$_5$ ($D_5$; [0-5 d]; $N_5$; 1.0 µM [5-10 d]) at 10 d. The bars and error bars indicate Mean ± SD. Significance was determined by one-way ANOVA using multiple comparisons test; ns = not significant.
